# Increased axon initial segment length results in increased Na^+^ currents in spinal motoneurones at symptom onset in the G127X SOD1 mouse model of Amyotrophic Lateral Sclerosis

**DOI:** 10.1101/2020.06.25.096552

**Authors:** H. S. Jørgensen, D.B. Jensen, K.P. Dimintiyanova, V.S. Bonnevie, A. Hedegaard, J. Lehnhoff, M. Moldovan, L. Grondahl, C.F. Meehan

**Author notes:** Contributed equally (D.B. Jensen, K.P. Dimintiyanova, V.S. Bonnevie).

## Abstract

Amyotrophic lateral sclerosis is a neurodegenerative disease preferentially affecting motoneurones. Transgenic mouse models have been used to investigate the role of abnormal motoneurone excitability in this disease. Whilst an increased excitability has repeatedly been demonstrated *in vitro* in neonatal and embryonic preparations from SOD1 mouse models, the results from the only studies to record *in vivo* from spinal motoneurones in adult SOD1 models have produced conflicting findings. Deficits in repetitive firing have been reported in G93A SOD1 mice but not in presymptomatic G127X SOD1 mice despite shorter motoneurone axon initial segments (AISs) in these mice.

These discrepancies may be due to the earlier disease onset and prolonged disease progression in G93A SOD1 mice with recordings potentially performed at a later sub-clinical stage of the disease in this mouse. To test this, and to explore how the evolution of excitability changes with symptom onset we performed *in vivo* intracellular recording and AIS labelling in G127X SOD1 mice immediately after symptom onset. No reductions in repetitive firing were observed showing that this is not a common feature across all ALS models. Immunohistochemistry for the Na^+^ channel Nav1.6 showed that motoneurone AISs increase in length in G127X SOD1 mice at symptom onset. Consistent with this, the rate of rise of AIS components of antidromic action potentials were significantly faster confirming that this increase in length represents an increase in AIS Na^+^ channels occurring at symptom onset in this model.

**Highights:**

- *In vivo* electrophysiological recordings were made in symptomatic G127X SOD1 mice.
- There were no deficits in repetitive firing in motoneurones in G127X mice.
- Increased persistent inward currents were still present in the symptomatic mice.
- Results suggest increases in Na^+^ currents at axon initial segments (AISs).
- Immunohistochemistry showed that motoneurone AISs were longer and thinner.

## INTRODUCTION

Amyotrophic lateral sclerosis (ALS) is a neurodegenerative disease, which preferentially affects motoneurones. Understanding why motoneurones, in particular, die in ALS is the holy grail of researchers in the field. One frequently investigated avenue is the excitotoxicity hypothesis and symptoms seen in humans such as spasticity and fasciculations (spontaneous activity from motoneurones) are indeed consistent with an increased excitability of lower motoneurones. The cell bodies of these neurones are relatively inaccessible in humans and so animal models of the disease provide a valuable tool to investigate spinal motoneurone excitability but have produced conflicting findings.

In a small subset of ALS patients, a number of different mutations have been identified and have led to the generation of transgenic mouse models which, to a variable extent, develop phenotypes consistent with the human disease. The most commonly used of these are the SOD1 mutants (Bruijn et al., 1997;Gurney et al., 1994;Jonsson et al., 2004;Jonsson et al., 2006;Ripps et al., 1995) which have been used extensively to study pathological alterations in the motoneurones that may contribute to their vulnerability in this disease, including excitability.

Experiments using cultured motoneurones or *in vitro* spinal cord slice preparations from neonatal SOD1 mutants have generally confirmed an increased intrinsic excitability of motoneurones. In particular, an increased gain in the current-frequency relationship and increased persistent inward currents (PICs), mediated by both Na^+^ and Ca^2+^ have been observed (Kuo et al., 2004;Kuo et al., 2005;Martin et al., 2013;Pambo-Pambo et al., 2009;Pieri et al., 2003;Quinlan et al., 2011;van Zundert et al., 2008) consistent with the hypothesis that an increased excitability may render the motoneurones vulnerable to excitotoxicity.

Disease onset in this disorder, however, is normally in adulthood. It is, therefore, crucial to determine whether the motoneurones still show an increased excitability in adulthood. Results from *in vivo* recordings in two different adult SOD1 mutants, however, have revealed partly consistent and partly conflicting findings. Recordings from motoneurones in adult presymptomatic G93A and G127X SOD1 mouse models have both shown generally normal excitability levels with respect to the input-output relationship (I-f slopes), voltage threshold for action potentials and rheobase (Delestree et al., 2014;Meehan et al., 2010). In the G93A SOD1 mice, however, a large proportion of motoneurones were reported as lacking the ability to fire repetitively which has been taken to indicate a hypo-excitability of some motoneurones (Delestree et al., 2014;Martinez-Silva et al., 2018). In the G127X SOD1 mouse no deficits in repetitive firing were observed, despite a slight decrease in the length of the axon initial segment (AIS), the action potential generating region of the neurone (Bonnevie et al., 2020). If anything, an increased excitability is suggested in the G127X SOD1 model by the indication of increases in persistent inwards currents (Bonnevie et al., 2020;Meehan et al., 2010).

One potential explanation for the discrepancies could be differences in the timing of disease onset and progression between the two models. In the G93A SOD1 mutant disease onset occurs around 90-120 days with a slow progression of around 3 weeks (Gurney et al., 1994). In the G127X SOD1 mouse, however, disease onset has been reported to occur much later at around 250 days followed by a rapid progression of approximately one week (Moldovan et al., 2012). It is therefore possible that the cells unable to fire repetitively in the G93A SOD1 model represent cells at a more advanced disease stage. Therefore, to investigate if motoneurone excitability is altered as symptoms emerge, we have recorded intracellularly from motoneurones again in the G127X mouse but now at symptom onset. We combined these functional measures with immunohistochemistry to investigate possible further reductions in the AIS length which has been shown to have consequences for repetitive firing (Evans et al., 2015). Surprisingly, we observed the opposite result with an increase in AIS length with detectable physiological consequences to action potentials. This data provides the first direct recordings of motoneurone excitability *in vivo* during the symptomatic stage in an ALS model and provides evidence for an increase in Na^+^ currents and/or channels at the AIS.

## METHODS

The experimental procedures described were approved by the Danish Animal Experiments Inspectorate (2012-15-2934-00501 and 2013-15-2935-00879) and were in accordance with the EU Directive 2010/63/EU. The methods described below were identical to those previously described in our publication in the same model at pre-symptomatic time points (Bonnevie et al., 2020;Meehan et al., 2010).

### G127X SOD1 Mice

The transgenic ALS mouse models used for this study expressed G127insTGGG (G127X) mutant human SOD1 and had been backcrossed on C57BL/6J mice for more than 25 generations in Umea, Sweden (Jonsson et al., 2004). Homozygote mice from the original line 716 which overexpresses 19 copies of the human SOD1 G127X were then bred as homozygotes at our own institution. This mutant SOD1 itself lacks SOD1 enzyme activity and is rapidly degraded. Consequently, low levels of the protein aggregate in the spinal cord and axons (Jonsson *et al.*, 2004). This all reduces possible overexpression artefacts which have been associated with the more commonly used G93A SOD1 mice. Experiments were performed on symptomatic mice, with the symptomatic stage defined as a slowing of movement, a hunched gait and extreme weakness or paralysis in at least one limb. This is roughly equivalent to a score of 3 on the ALSTDI scoring system for ALS mutants (Hatzipetros et al., 2015). As the mutant SOD1 G127X has no enzymatic function and for technical reasons, it was necessary to breed the G127X mice as homozygotes the best available controls that could be used were roughly aged-matched C57BL/6J mice (the background strain for the G127X mice) which we will refer to as wild type (WT). For the electrophysiological comparisons between WT and symptomatic G127X mice we used the data from the same WT mice as in our previous study (Bonnevie et al., 2020) where the mice were only ~1 month younger than here. New WT mice were, however, used for the AIS labelling, as the same reagents (in particular, the same lot number of the polyclonal antibody) were not available as used in the previous study. The use and details of the mice used for the different parts of the investigation are provided in Table 1.

**Table1.** Details of the mice used for the different parts of the investigation. In accordance with ethical principles (the 3 Rs), 3 of the G127X mice were also used for experiments on post-activation depression (Hedegaard et al., 2015) and the mice used for the electrophysiology experiments were also used for investigating axonal potassium channel distribution and function (Maglemose et al., 2017).

|  | Group | N (mice) | Male/Female | Mean Age (days) | Age Range (days) |
| --- | --- | --- | --- | --- | --- |
| AIS labelling | WT | 9 | 4/5 | 246 | 217-250 |
|  | G127X | 6 | 5/1 | 226 | 209-235 |
| Cell counting | WT | 5 | 0/5 | 220 | 215-240 |
|  | G127X | 5 | 0/5 | 223 | 212-240 |
| Electrophysiology | WT | 6 | 0/6 | 193 | 182-210 |
|  | G127X | 7 | 4/3 | 232 | 212-281 |

### Electrophysiological Experiments

*In vivo* electrophysiological experiments were performed precisely as described in our previous studies (Bonnevie et al., 2020;Meehan et al., 2010). Anaesthesia was initially induced (under brief isoflurane exposure to reduce stress) and maintained with IP injections of Hypnorm (fentanyl-citrate 0.315mg/ml) and Midazolam (fluanisone 10 mg/ml) in dH_2_O (mixed 1:1:2, induction dose 10ml/kg, supplemented with 0.05ml every 20 minutes). A single dose of atropine (0.02mg IP) was administered at the start to reduce mucous secretion before a plastic cannula was inserted into the trachea and IP cannulae inserted and secured in place for further drug administration. After unilateral dissection of the sciatic nerve (on the left side) a hemi-laminectomy (also left side) was performed at vertebral levels T12-L1. This provided access to spinal segments L3-L4 which contains the majority of motoneurones with axons in the sciatic nerve in mice.

Mice were placed in a modified Narishige stereotactic frame. Here, the temperature was monitored using a rectal probe and was maintained at 37°C by a heating pad beneath the mouse and a heat lamp above, both controlled by a system using feedback from the temperature probe. The tracheal cannula was connected to a ventilator (SAR-83 CWE) to allow the mice to be artificially ventilated at ~70 breaths per minute (with a tidal volume of approximately 0.2ml). Once ventilated, the neuromuscular blocking agent Pavulon was administered (diluted 1:10 with saline then 0.1ml dose initially followed by 0.05ml doses every hour). Anaesthesia was maintained using the same dosages as previously required for the surgical procedures. The electrocardiogram was monitored using clips placed on the ear and rear foot to monitor anaesthesia under neuromuscular blockade and expired carbon dioxide levels were also measured using a Capstar CO_2_ analyser (IITC Life Science).

In the recording frame, Narashige vertebral clamps were used to stabilise the vertebrae immediately above and below the laminectomy site to stabilise and the sciatic nerve was mounted on custom-made hook electrodes for antidromic stimulation. An electronic micro-drive was used to drive a glass microelectrode (filled with 2M potassium acetate, pipette resistance ~27 mΩ) into the spinal cord and antidromic field potentials from stimulation of the peripheral nerves were used to locate the hind limb motoneurones. Signals were amplified and filtered (10kHz) using custom made amplifiers. The amplified signals were digitised (at 40kHz for intracellular recordings) using the 1401 analogue to digital converter (Cambridge Electronic Design, UK) and recorded and analysed using Spike 2 software (Cambridge Electronic Design, UK).

Custom-made silver ball electrodes placed on the dorsolateral surface of the exposed spinal cord recorded the incoming volley for sensory input to the spinal cord from the peripheral nerve stimulation. The timing of this cord dorsum potential was used to distinguish between antidromic (back-fired) and synaptically evoked action potentials, allowing antidromic identification of the motoneurones. Antidromic action potentials from successive stimuli were averaged for analysis and the resting membrane potential (Vm) was measured at the point at which an average was made (confirmed using extracellular potentials upon exiting the cell dorsally). All analysis of spike height was performed taking resting Vm into account with action potentials excluded from analysis if the Vm could not be reliably confirmed, if the Vm was more depolarized than −50mV or if there was no overshoot of the action potential. The average of the action potential was differentiated (dV/dt) using Microsoft Excel and the maximum rate of rise of the initial segment (IS) and soma dendritic (SD) components were determined from the first derivative. The time difference between these peaks representing the IS and SD components was used to measure the IS-SD delay (Gustafsson and Lipski, 1980).

To test the ability of the motoneurones to fire repetitively in response to central inputs, intracellular current injections were used to mimic synaptic input. Triangular current ramps were injected through the microelectrodes using DDC mode (3 kHz) on the Axoclamp amplifier and the cell’s ability to fire repetitively determined. Special care was taken to confirm that the electrode capacitance was correctly compensated (using the capacitance compensation on the Axoclamp 2b amplifier) and that the electrode was passing the current effectively (important given the high resistance of the electrodes used for such experiments). In addition, the membrane potential at the time point immediately prior to current injection was also confirmed to be more hyperpolarised than −50 mV as this indicates a reliable penetration (important for repetitive firing). The current-frequency (I-f) gain was obtained during the primary range (Granit et al., 1966a) and measured using the triangular ramps with instantaneous firing frequencies computed in Spike-2 software and then exported to Microsoft Excel. Where possible, the Na^+^ PIC was also estimated from the triangular ramps by subtraction of the actual rheobase current during the triangular current from the estimated rheobase current if the voltage response to the current injection was linear (as in (Delestree et al., 2014)).

Input resistance was calculated using the voltage drop measured in response to a 3nA (~50ms) hyperpolarising current delivered through the microelectrode (3-5 kHz). Analyses were performed using averages of multiple trials. To measure the effects of hyperpolarization-activated (Ih) currents the sag was measured (difference in voltage at the start and end of the 50 ms current pulse) and expressed as both absolute voltage sag and as a percentage of the absolute voltage drop. The Vm immediately after the removal of the hyperpolarising current was measured and the difference between this and the resting Vm was calculated to determine the rebound.

### Anatomical Experiments

Protocols for the anatomical experiments were also identical to those used in the previous pre-symptomatic study (Bonnevie et al., 2020). After electrophysiological experiments mice were perfused via the left ventricle with saline followed by a 5-minute perfusion with 2% paraformaldehyde (diluted in phosphate buffered saline, pH7.4). The remaining vertebrae were removed and spinal cords were removed and post-fixed for 1 hour in 2% paraformaldehyde and further stored overnight in 30% sucrose for cryo-protection during cutting. The lumbar enlargement was cut into 50 µm thick horizontal sections on a freezing microtome (HM450, Thermo Scientific) and sections from the ventral portion of the spinal cord containing motoneurones were selected for processing.

Axon initial segments were labelled with a primary antibody against the main voltage-gated Na^+^ channel found at the AIS of motoneurones responsible for action potential initiation; Nav 1.6 (rabbit anti-Nav 1.6, Alomone, catalogue number: ASC-009, 1:500). This Nav 1.6 antibody produced the expected pattern consistent with axon initial segments and nodes of Ranvier. Secondary-only controls and pre-incubation with the immunizing peptide resulted in no labelling (not shown). The motoneurones were labelled with an antibody directed against Choline acetyltransferase (goat anti-ChAT, Millipore, catalogue number: AB144P, 1:100). This antibody labelled known cholinergic populations of neurones in the correct locations including large ventral horn neurones (motoneurones), sympathetic preganglionic neurons in the intermediolateral nucleus (IML) and small cholinergic interneurones around the central canal and intercalated nucleus. Primary antibodies were visualised using secondary antibodies conjugated to the Alexa Fluor fluorescent dyes 594 and 488 (Invitrogen/Thermofisher catalogue numbers: A-11055 and A-21207).

#### Measurements of the AIS

Anatomical analyses of AISs were performed in 3-D using z-stacks acquired with a LSM700 Zeiss confocal microscope and analysed using Zen software (Zeiss). The length and width (at the proximal end which is normally the widest part) of the AISs was measured as well as the distance from the proximal end of the AIS to the soma and the 2D area of the soma (calculated as the area of an ellipse obtained from the minimum and maximum diameters). Gamma motoneurones were classified by a size criteria (less than 485 μm^2^ (Shneider et al., 2009)) and a lack of cholinergic bouton labelling around the soma. To be included the AIS had to connect to a ChAT-immunoreactive cell and the entire AIS had to be contained within the confocal stack and not be cut-off neither in the z-nor in the xy-plane (confirmed by at least one confocal slice above and below the AIS containing labelling of other processes). For measurements of distance from the soma, AISs which originated from dendrites (as occurs occasionally on spinal motoneurones) were excluded from analysis.

### Cell counting

An additional 10 mice (5 WT and 5 G127X) were used for cell counting. These G127X mice were all symptomatic with similar ages to those used for the rest of the experiments (Table 1). Serial transverse 40 μm thick sections were cut from the lower thoracic region to the upper sacral part of the spinal cord. From these, every 10^th^ section was processed immunohistochemically for ChAT using the same antibody as above and a donkey anti-goat Alexa 594 secondary antibody (Invitrogen/Thermofisher catalogue numbers: A-11058). Motoneurone counts were made manually using an epifluorescent microscope (Axioplan 2, Zeiss, Germany). Only motoneurones with visible nuclei were counted. The rostral-caudal extent of the lumbar region of the spinal cord for analysis was defined by the end of the IML (rostral border) and start of the sacral parasympathetic (SPSy) nucleus (caudal border) which mark the first lumbar segment and the beginning of the sacral segments, respectively. The number of motoneurone per section were plotted in serial order from rostral to caudal and aligned with respect to these borders and the peak of the distribution. For 5 G127X mice (3 of the above and 2 additional) double labelling was also performed (on every 20^th^ section) using antibodies against ChAT (as above) and the apoptotic marker cleaved caspase-3 (rabbit anti-cleaved Caspase-3 (Asp175) (5A1E), Cell Signalling catalogue number: 9664 1:500) visualised using the following secondary antibodies conjugated to fluorochromes; donkey anti-goat Alexa Fluor 488 (1:1000) and donkey anti-rabbit Alexa Fluor 594 (1:500) respectively (both from Invitrogen).

Illustrative photographs of immunohistochemical labelling were obtained with the same confocal microscope or epifluorescent microscope as described above. Brightness and contrast were adjusted using Image J (NIH) or Photoshop (Adobe) and applied uniformly to the entire image for illustration purposes only. All analyses, however, were performed with the raw data.

### Statistical analysis

All statistical tests were performed using the GraphPad Prism software. D'Agostino & Pearson omnibus normality tests were used to confirm normality and parametric (un-paired t-tests) or non-parametric (Mann-Whitney) tests were used accordingly. In the case of a low sample size, non-parametric statistics were automatically used. Linear regression was performed for electrophysiological parameters affected by Vm and then the slopes were tested for significant difference between either the angle of the slopes or the elevation (intercept). Statistical significance was accepted at the P≤ 0.05 level. On all graphs stars are used to indicate the following significant differences: * (P ≤ 0.05), ** (P ≤ 0.01), *** (P ≤ 0.001), **** (P ≤ 0.0001). Unless indicated on the graphs no significant difference was found.

## RESULTS

### AISs of alpha motoneurones are longer and thinner in G127X mice

Examples of AISs labelled with antibodies against Nav1.6 are shown in Fig. 1A. The measurements of the different parameters for alpha motoneurones, along with more details of the statistical analyses are shown in Table 2 and illustrated as scatterplots in Fig. 1. Axon initial segments of alpha motoneurones were 12.9% longer in the G127X mice than in WT mice (P<0.0001, Fig. 1B). When displayed as cumulative frequency plots it can be observed that the entire population of AISs are shifted towards longer lengths (Fig. 1C). AISs were also slightly thinner proximally by 6.3% (P=0.0382, Fig. 1D) but no significant differences were found with respect to distance of the proximal end of AIS to the soma (Fig. 1E). The cell bodies of the motoneurones in the G127X mice were 11.6% larger than WT (P=0.0013, Fig. 1F).

**Figure 1.**
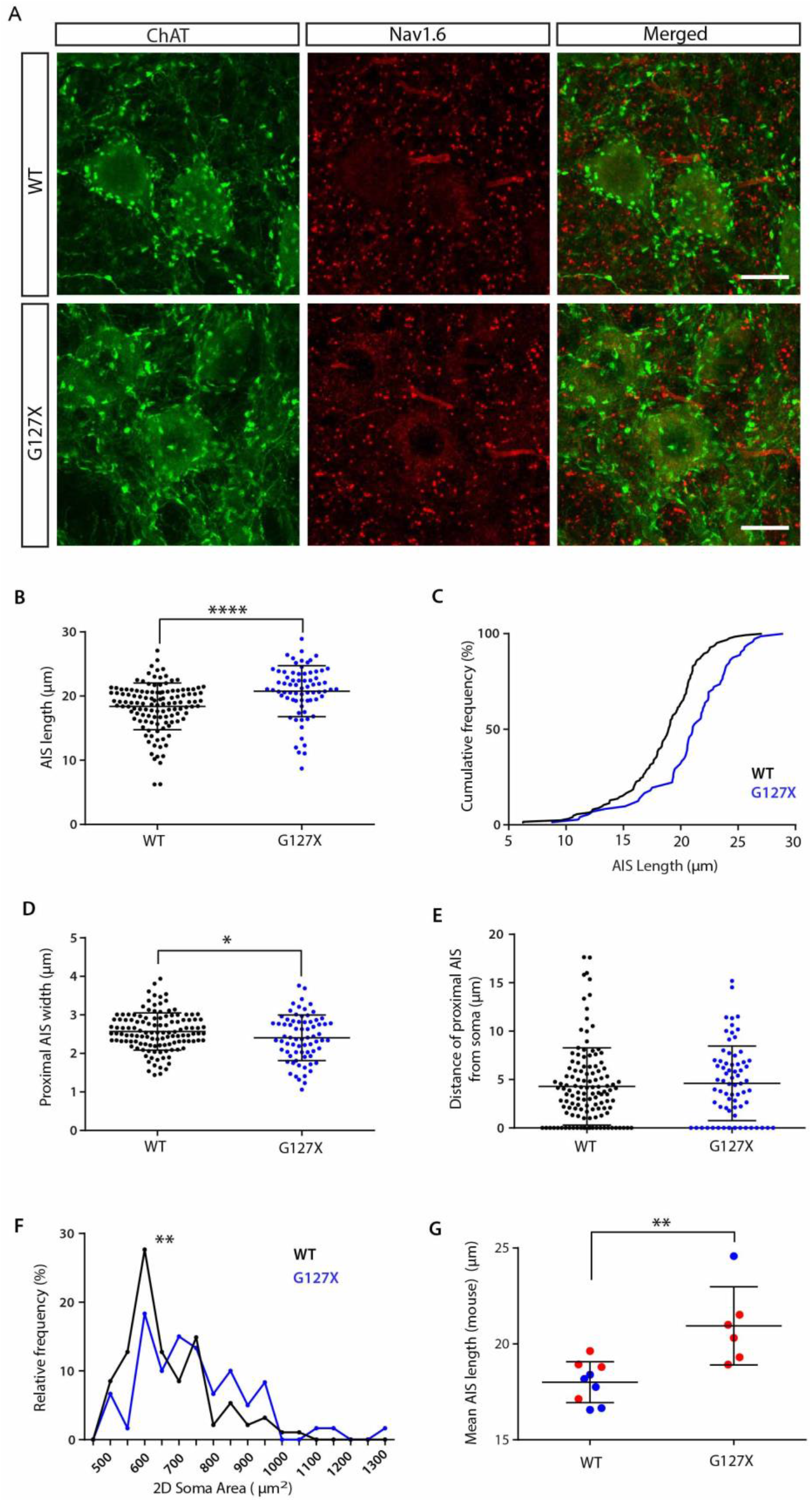
AIS parameters of alpha motoneurones in WT (black) and G127X (blue) mice. A. Immunohistochemical labelling of Nav 1.6 (red) at AISs and ChAT (green) immunoreactive alpha motoneurones from WT and G127X mice. Scale bar 20μm. B. Scatter dot plot showing values for individual cells for AIS length of alpha motoneurones, which are significantly longer in the G127X mice (P < 0.0001, n= 124 WT cells, 72 G127X cells). C. Cumulative frequency distribution of AIS length of alpha motoneurones showing a shift of the entire distribution towards longer AIS lengths in G127X mice. D. Scatter dot plot of values for individual cells for the proximal AIS width of alpha motoneurones. This shows a significant reduction in AIS width in the G127X mice (P= 0.0382, n= 125 WT cells, 72 G127X cells). E. Scatter dot plot showing values for individual cells for the distance of the proximal AIS from the soma in alpha motoneurones, which was not significantly different (n= 125 WT cells, 72 G127X cells). F. Histogram showing the frequency distribution of 2D soma area of alpha motoneurones grouped in 50μm bins. From this it can be seen that soma size is larger in the G127X mice (P= 0.0013, n= 94 WT cells, 60 G127X cells). G. Scatter dot plot showing the mean AIS length for each animal with the mean value for each animal represented as a single dot (red indicates male, blue indicates female, P= 0.0028, n= 9 WT mice, 6 G127X mice). In all scatter dot plots, the lines show overall means and standard deviations.

**Table 2.** AIS parameters with mixed sex groups.

|  | WT |  | G127X |  | Significance |
| --- | --- | --- | --- | --- | --- |
|  | Mean (SD) | N (cells) | Mean (SD) | N (cells) |  |
| Length ($\mu\text{m}$) | 18.39 (3.639) | 124 | 20.76 (3.966) | 72 | **P < 0.0001** |
| Distance from soma ($\mu\text{m}$) | 4.287 (3.994) | 125 | 4.601 (3.849) | 72 | P = 0.3933 |
| Width ($\mu\text{m}$) | 2.569 (0.4836) | 125 | 2.406 (0.5933) | 72 | **P = 0.0382** |
| 2D Surface Area ($\mu\text{m}$) | 665.9 (122.6) | 94 | 743.1 (159.6) | 60 | **P = 0.0013** |

Given that the WT and G127X groups consisted of uneven proportions of males and females we first compared male and female within the WT group to see if there were any general sex differences in WT mice. There were no significant differences between AISs of motoneurones from WT males and females with respect to any of the AIS parameters measured (length, width or distance from soma). There was, however, a significant difference between males and females with respect to cell size, with motoneurones from the males being significantly larger than the females (P=0.0024).

To determine whether the differences in AIS length and cell size that we observed in the G127X mice was due to a larger proportion of males (with larger somas) we repeated our analysis but only including males in both groups. The data is given in Supplemental Table 1. The same patterns were observed in the male only group as before with significantly longer and thinner AISs in the G127X males compared to the WT males. There was also still a significant increase in soma size in the male G127X mice compared to the male WT mice. The distance of the proximal AIS from the soma was still not significantly different from one another. To test if the increase in AIS length was consistent across all mice the mean AIS length was calculated for each mouse, including both males and females (illustrated in Fig. 1G). Here, a significant difference was still observed, in fact, when analysed this way, the mean length was 16.9% longer in the G127X mice (P= 0.0028, Mann Whitney). The decrease in AIS width and increase in cell size were not significantly different when compared using means per mice (P= 0.1726 and P=0.7039 respectively) although we must express caution with these particular analyses conducted on a mean per animal basis, due to the low numbers of mice and high variability within each parameter (apparent on Fig.1). Finally, to confirm that the increased AIS length was not due to larger cells in the G127X mice the relationship between the two parameters was tested for cells with the entire soma contained within the z-stack. Linear regression showed that, in this sample, there was not a significant relationship between AIS length and soma size for either WT (R^2^=0.009115, P= 0.3626 n= 93 cells) or G127X (R^2^= 0.03379, P= 0.1598, n=60 cells), if anything, the smaller cells in the G127X mice tended to the have longer AISs.

Gamma motoneurones were identified by size as described in the methods and excluded from the initial analyses of alpha motoneurones. As a separate group there was a visible but not statistically significant tendency (P=0.0684) for AISs of gamma motoneurones to also be longer by approximately 11.4% in G127X mice (supplementary table 2). However, a low number of cells ended up in this group (34 WT and 21 G127X across all mice) precluding a more rigorous analysis of these neurones as an independent group and so further experiments would be needed to confirm this tendency.

### No change in somatic action potential amplitude in G127X motoneurones

To investigate whether the changes in AIS parameters have a functional effect, intracellular recordings were performed. Examples of averaged antidromic action potentials from both WT and G127X motoneurones along with the first derivatives (dV/dt) of the averages are shown in Fig. 2A and B. Antidromic action potentials were recorded from the soma of 126 motoneurones (89 WT and 37 G127X) and had the characteristic shape consisting of an inflection on the upward phase due to the presence of the action potential both when it arrives at the AIS and, after a short delay when it arrives at the soma (seen as the peak). The IS (initial segment spike) and SD (soma-dendritic spike) components can be clearly appreciated from the first derivative (Fig. 2A, B) with two positive peaks representing the maximum rate of rise of the IS and SD components, respectively (Lipski, 1981).

**Figure 2.**
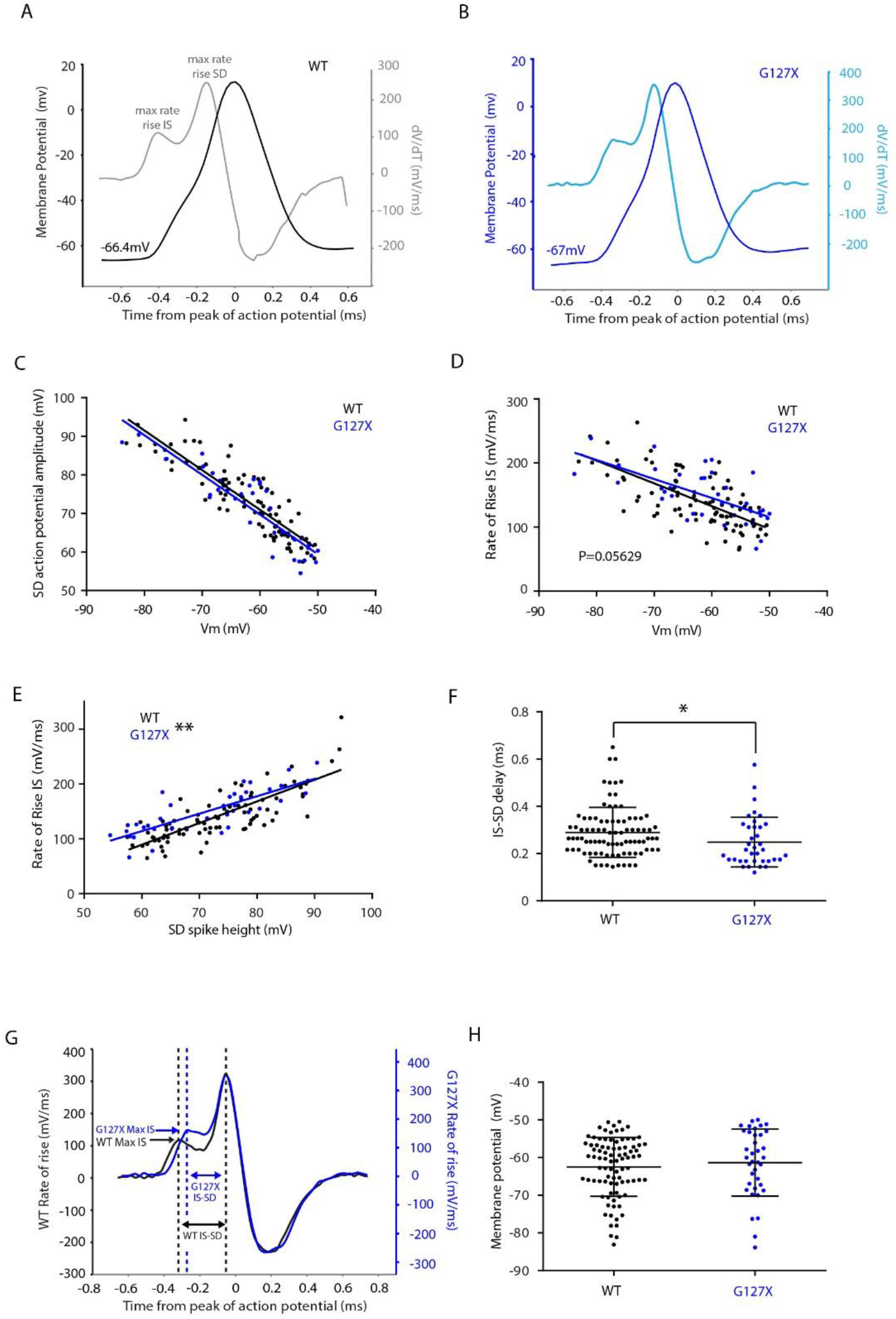
Features of antidromic action potentials of motoneurones in WT (black) and 37 G127X (blue) mice. A. Example of an average of antidromic action potentials (black) from a motoneurone in a WT mouse with its first derivative (grey) showing two peaks corresponding to the maximal rate of rise of the IS and SD components of the antidromic action potential. B. Example of an average of antidromic action potentials from a motoneurone in a G127X mouse (dark blue) with its first derivative (light blue). C. Data points for individual action potentials and linear regression lines showing the significant relationship between SD action potential amplitude and resting Vm for motoneurones from WT and G127X mice. There was no significant difference between these two regression lines with respect to both slope and elevation. D. Data points for individual action potentials and linear regression lines showing the relationship between the maximum rate of rise of the IS component of the action potential and Vm for motoneurones from WT (black) and G127X (blue) mice. From this, a tendency for the rate of rise of the IS spike to be faster in the G127X mice can be seen between the two slopes with respect to elevation (P=0.05629). E. Data points for individual action potentials and linear regression lines showing the relationship between the amplitude of the maximum rate of rise of the IS component of the antidromic action potential and SD spike height from WT (black) and G127X (blue) mice. Here a significant difference between the slopes is seen (P= 0.00318) with respect to elevation, consistent with an increase in AIS Na^+^ currents relative to somatic Na^+^ currents. F. Scatter dot plot showing data points for individual cells for IS-SD latency for motoneurones, measured as the time interval between the maximum rate of rise of the IS and the SD components of the antidromic action potential. Lines show means (and SD) which was significantly faster in the G127X mice (P= 0.0172). G. Representative examples of first derivatives of antidromic action potentials from WT and G127X mice illustrating the faster rate of rise of the IS component relative to the SD component and the shorter IS-SD delay in G127X mice. H. Scatter dot plot showing no significant difference in the resting Vm for the motoneurones at the time point at which the antidromic action potentials were tested. Lines show means (and SD). (*: P < 0.05; **: P < = 0.01, n= 89 WT and 37 G127X cells)

The maximum amplitude of the antidromic action potentials (SD component) correlated significantly with resting Vm for both WT (R^2^=0.7788) and G127X motoneurones (R^2^=0.8136, both P< 0.0001, Fig. 2C). The regression slopes were compared and no significant differences were found between WT and G127X both with respect to the slope (P= 0.9971) and the elevation (P= 0.1522, Fig. 2C). The maximum rate of rise of the SD component of the antidromic action potential also correlated significantly with the resting Vm for both WT (R^2^=0.3283) and G127X (R^2^= 0.4638) motoneurones (both P< 0.0001). Regression slopes comparisons revealed no significant differences between WT and G127X both with respect to the slope (P= 0.1524) and the elevation (P= 0.6226). Together, these results suggest no changes in somatic Na^+^ currents in the G127X mice.

### Initial segment components of antidromic spikes have a faster maximum rate of rise

The maximum rate of rise of the IS component of the antidromic action potential was also confirmed to correlate significantly with somatic Vm by using linear regression for both WT (R^2^=0.4488) and G127X (R^2^=0.4315) motoneurones (both P< 0.0001 Fig. 2D). The regression slopes were compared and no significant difference was observed with respect to the slope (P= 0.4172); however, an observable difference was observed with respect to the elevation with the maximum rate of rise of the IS component being faster for G127X mice but this just failed to reach statistical significance at the P<0.05 level (P=0.05629). These results suggest that the relationship between maximum rate of rise of the IS component and Vm is altered in the G127X motoneurones in favour of increased Na^+^ currents at the AIS. This was confirmed by linear regression between the spike height of the full antidromic action potential (i.e. the amplitude of the SD component) and the maximum rate of rise of the IS (Fig. 2E). This relationship was significant for both WT (R^2^=0.6564) and G127X (R^2^=0.6431) groups at the P<0.0001 level. Regression slopes were compared between the groups and no significant difference was observed with respect to the slope (P= 0.145), however, a significant difference was observed with respect to the elevation, with the maximum rate of rise of the IS component being faster for G127X mice (P=0.003178). This confirmed that the maximum rate of rise of the IS component in the G127X mice is proportionally faster than WT for the same SD action potential height.

A faster rate of rise of the IS component may be expected to depolarise the surrounding area faster and may lead to a faster propagation of the action potential from the AIS to the soma. To test this hypothesis, we used the time difference between the maximum rate of rise of the IS and the SD components of the antidromic action potential on the dV/dt plot as a measure of the IS-SD delay. Consistent with this hypothesis, the IS-SD delay was significantly shorter in the G127X mice (means (+SD): WT: 0.29 ms (0.105), G127X: 0.25ms (0.105), P=0.0172, Fig. 2F). Examples of two dV/dt curves representative of the average features for both groups are shown in Fig. 2G illustrating the differences observed. To rule out whether the reduction in IS-SD delay was due to the soma being more depolarised in the G127X we compared the resting Vm immediately before the action potentials for the two groups. There was no significant difference in Vm between the groups (means (+SD): WT: −62.48 mV (7.8), G127X: −61.43 mV (8.89), P=0.3795 Fig. 2H), therefore, this cannot explain the reduction in IS-SD delay observed in the G127X mice. Also, a decreased distance of the AIS from the soma was not observed in our anatomical experiments eliminating this possible explanation.

### Repetitive firing is not impaired in the G127X motoneurones

Triangular current ramps evoked repetitive firing in 94.6% (53/56) motoneurones in WT mice and 90% (27/30) motoneurones in G127X mice (Fig.3 A, B). The slight reduction in the proportion of cells firing was not statistically significant (P=0.405, Fisher’s exact test) and would equate to a single cell difference (Fig. 3B). The features of the repetitive firing were measured, as illustrated in Fig. 3C, in those cells with stable firing. The voltage threshold and rheobase were measured using these ramps. Rheobase currents, appeared, on average, to be smaller in G127X mice (Fig. 3D) but statistical significance was not reached (means (+SD): WT: 7.18nA (3.77) G127X: 5.7nA (3.38), P=0.0974, n= 49 WT cells and 26 G127X cells). Similarly, there was an appreciable, but not significant tendency for the voltage threshold in motoneurones from G127X mice to be more hyperpolarised than in WT (means (+SD): WT: −48.32 mV (4.35) 44 cells, G127X: −50.62 mV (5.08) 25 cells, P=0.0507, Fig. 3E).

**Figure 3.**
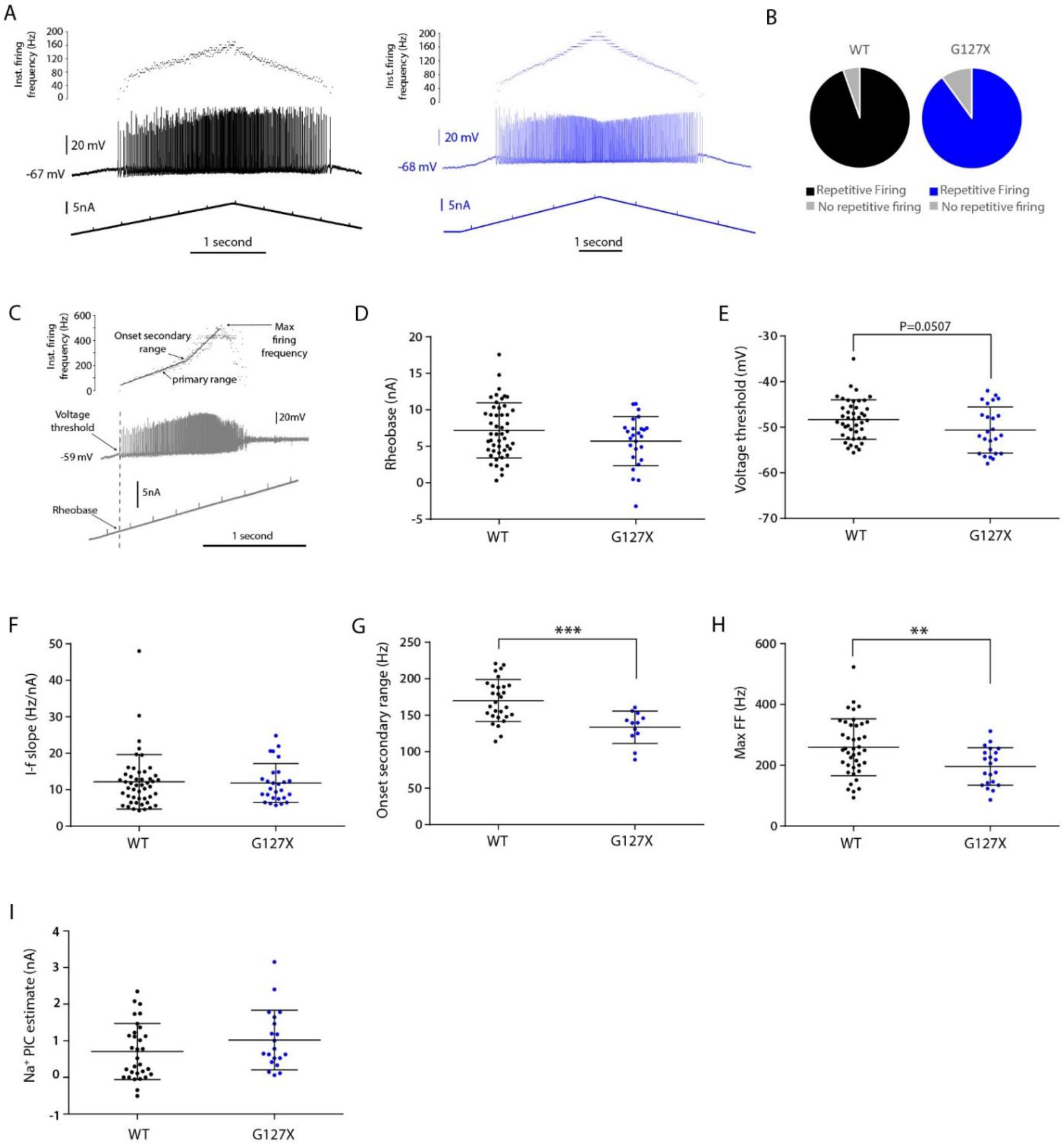
Repetitive firing in motoneurones in response to intracellular current injection in WT (black) and G127X (blue) mice. A. Example of repetitive firing recorded intracellularly (middle traces) in motoneurones from a WT and a G127X mouse in response to intracellular injection of ramps of ascending and descending intensity of current (bottom trace). The instantaneous firing frequency is shown on the top trace. B. The proportions of motoneurones in WT and G127X mice, responding to the current injection with repetitive firing which was not significantly different (P= 0.405, n= 56 WT cells and 30 G127X cells). C. An example of repetitive firing in a motoneurone in which the current has been taken up to the level where repetitive firing fails to determine the maximum firing frequency. An inflection can be seen in the I-f slope corresponding to the transition from the primary to the secondary range of firing. The way in which the voltage threshold and rheobase were determined from this is also illustrated. D. Scatter dot plot showing data points for individual cells for the rheobase currents at which repetitive firing was evoked in motoneurones, which, on average, was lower in G127X mice but not significantly different (P= 0.0974, n= 49 WT cells and 26 G127X cells). E. Scatter dot plot showing data points for individual cells for the voltage thresholds at which repetitive firing was evoked in motoneurones, which tended towards being at more hyperpolarized values in the G127X mice (P= 0.0507, n= 44 WT cells and 25 G127X cells). F. Scatter dot plot showing data points for individual cells for I-f slopes in the primary range, which was not significantly different (P= 0.9498 n= 48 WT cells and 27 G127X cells). G. Scattered dot plot showing data points for individual cells for the firing frequencies at which the transition from the primary range to the secondary range occurs, showing that the secondary range has a significantly earlier onset in the G127X mice (P= 0.0003, n= 29 WT cells and 12 G127X cells). H. Scatter dot plot showing data points for individual cells for the maximum firing frequencies of motoneurones showing a significant reduction in motoneurones with high maximum firing frequencies in G127X mice (n= 41 WT cells and 21 G127X cells). I. Scatter dot plot showing individual data points for the Na^+^ PIC estimated by subtraction of the actual rheobase current during the triangular current ramp from the estimated rheobase current if the voltage response to the current injection was linear. A non-significant tendency towards this being higher in G127X mice is seen (P= 0.1687, n= 31 WT cells and 20 G127X cells). In all scatter dot plots the lines show means and standard deviations.

When the current-frequency (I-f) slopes were calculated using the primary range of firing, no significant difference was observed between WT and G127X (means (+SD), WT: 12.18 Hz/nA (7.48) n= 48 cells, G127X: 11.82Hz/nA (5.36) n= 27 cells, P=0.9498, Fig. 3F). After the primary range, in some cells, a steep inflection can be seen (the secondary range, Fig. 3C) as a result of the activation of dendritic persistent inward currents (Granit et al., 1966b). The onset of the secondary range occurred at significantly lower firing frequencies in G127X mice (means (+SD): WT: 170.2 Hz (+28.9), 29 cells, G127X: 133.6 Hz (22.17) 12 cells, P=0.0003, Fig. 3G).

In 62 of the motoneurones the current injection was taken to the level where repetitive firing of full spikes failed in order to determine the maximum firing frequencies in response to the ramp current injection. Here, a significant reduction in maximum firing frequencies was observed in the G127X mice (means (+SD): WT 259 Hz (93.47) n= 41 cells, G127X 196 Hz (61.94) n= 21 cells. P= 0.0071, Fig. 3H). From Fig. 3H it appears that the overall range of maximum firing frequencies is reduced in the G127X mice and is consistent with of a loss of cells with higher maximum firing frequencies. F tests suggested that the standard deviations of the 2 groups were different and bordered on significance (P=0.0512).

Due to the presence of mixed-mode oscillations (Manuel et al., 2009), synaptic noise and a low DCC switching frequency it was only possible to estimate the Na^+^ PIC in 31 WT and 20 G127X motoneurones (Fig. 3I). The mean value was higher in G127X mice but not significantly different (means (+SD): WT: 0.71 nA (0.77), G127X: 1.02 nA (0.82), P= 0.1697).

### Evidence for increased Ih currents in G127X motoneurones

To test whether the increase in cell size that we measured was associated with a decrease in input resistance this was measured using the voltage response to a brief (~50 ms) 3 nA hyperpolarizing current pulse (Fig. 4A). No significant difference was observed with respect to input resistance (means (+SD): WT 4.41 MΩ (2.59), G127X, 3.70 MΩ (1.71), P= 0.3779, Fig. 4B). After the initial voltage drop there was often a sag representing the activation of an I_h_ current from hyperpolarization-activated (HCN) channels. This occurred more frequently in the G127X mice (WT: 25/49 cells, G127X: 18/21 cells, Fishers exact test, P=0.0074). Where present, this sag was measured and was significantly more pronounced in the G127X mice, both if expressed as absolute voltage sag (means (+SD): WT: 1.267 mV (0.761), G127X: 1.996 mV (1.146), P=0.0166, Fig. 4C) or as a percentage of the total voltage drop to control for differences in input resistance (means (+SD): WT: 11.01% (5.541), G127X: 20.56% (10.51), P=0.0004, Fig. 4D). The increased activation of HCN channels resulted in an increased rebound with a larger overshooting of resting Vm upon removal of the hyperpolarising current (means (+SD): WT: 1.357 mV (0.7131), G127X: 1.857 mV (0.8471), P=0.0443, Fig. 4E). These results are therefore consistent with increased Ih currents in the G127X mice.

**Figure 4.**
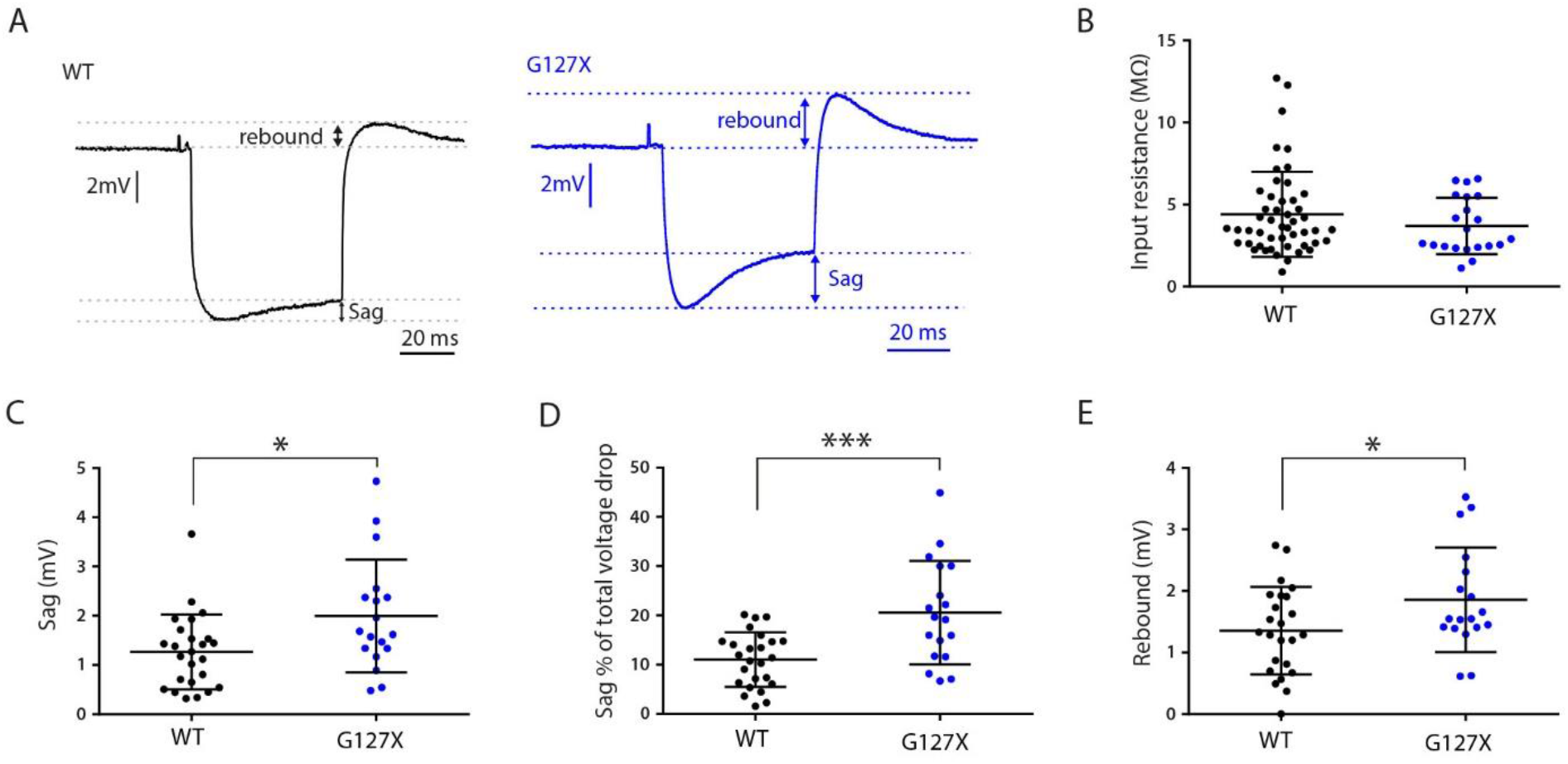
Input resistance and sag properties of motoneurones from WT (black) and G127X (blue) mice. A. Examples of the voltage drop in response to intracellular injections a 50ms, 3nA hyperpolarising square current pulses from motoneurones in WT and G127X mice. From this a depolarising “sag” in the response can be seen in many motoneurones, followed by a depolarising rebound response immediately after the end of the pulse. This was observed more frequently in the G127X mice (WT: 51%, n = 29 cells, G127X: 85.7%, n= 21 cells, P= 0.0074). B. Scatter dot plot showing the absolute input resistances for individual cells calculated from the peak voltage drop, which was not significantly different (P= 0.3779, n=49 WT cells, 21 cells). C. Scatter dot plot showing the sag recorded at the end of the 50ms long −3nA current pulse. This was measured in all cells showing a sag and was significantly larger magnitude in motoneurones from G127X mice (P= 0.0166, n=25 WT cells, n= 18 cells). D. Scatter dot plot for individual cells showing the same sag expressed as a percentage of the total voltage drop, which is also of larger magnitude in motoneurones from G127X mice (P= 0.0004, n=25 WT cells, 18 cells). E. Scattered dot plot for individual cells showing the rebound depolarisation observed after removal of the hyperpolarising current showing that this is also significantly enhanced in motoneurones in G127X mice (P= 0.0443, note, this was not possible in one cell due to post inhibitory rebound firing, n= 24 WT cells, n= 18 cells).

### Minimal motoneurone loss at symptom onset in G127X mice

To determine whether any of the differences we have observed were due to a selective cell loss we quantified cell loss in 5 symptomatic G127X and 5 age-matched WT mice. The mean number of motoneurones at every 10^th^ 40 μm thick transverse spinal cord section from rostral to caudal is shown in Fig. 5A. From this dataset it can be observed that although there appears to be cell loss (most obvious at the centre of the lumbar enlargement) it is surprisingly minimal given that the G127X mice were clearly symptomatic. The mean number of motoneurones per section across the lumbar cord was lower in G127X mice (P= 0.2460, Mann Whitney, means (+SD): WT: 53.2 (3.75), G127X: 48.63 (7.34)), but this was not significantly different (although this comparison has limited statistical power). A significant difference was only seen if specifically comparing the 7^th^ section, i.e. the very centre of the lumbar enlargement (P= 0.0079, Mann Whitney, means (+SD): WT: 58.60 (6.11), G127X: 41.20 (6.84)). Cleaved caspase-3 labelling, however, revealed large clusters of motoneurones actively undergoing the process of apoptosis in all 5 of the G127X mice tested (Fig. 5B). The labelling also revealed cleaved-caspase 3 activity in other populations of ChAT immunoreactive neurones including the small cholinergic neurones surrounding the central canal (Fig. 5C) and sympathetic preganglionic neurones (Fig. 5D) in the intermediolateral nucleus (in the most rostral sections tested). This phenomenon was not observed in WT mice.

**Figure 5.**
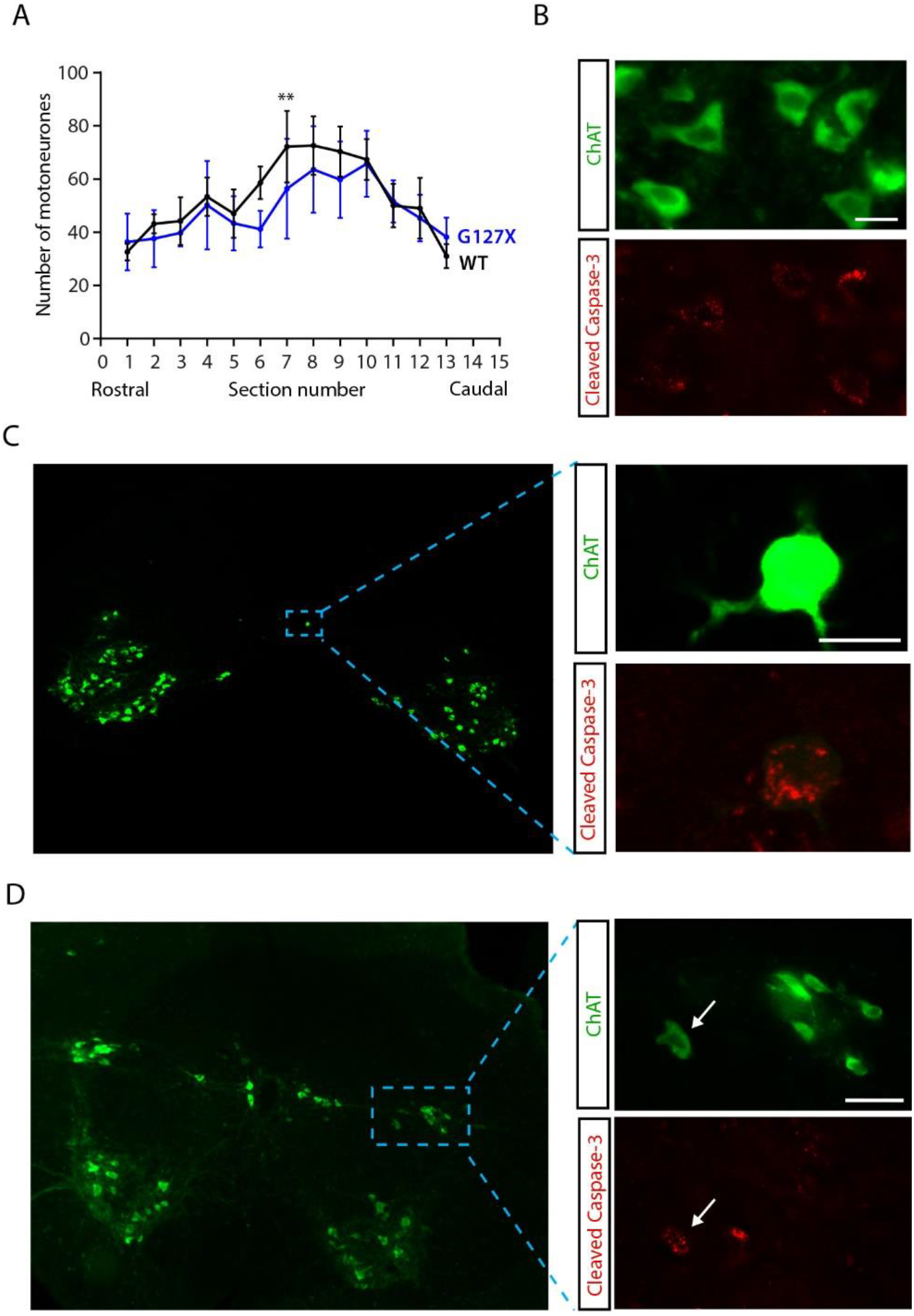
Motoneurone loss and cleaved caspase-3 activation in ChAT immunoreactive motoneurones, IML and central canal neurones in G127X mice. A. Frequency histogram showing the mean number of motoneurones counted per section in serial order from rostral to caudal sections in lumbar spinal cords of WT (black, n= 5) and G127X (blue, n= 5) mice. A loss of cells can be observed at the centre of the lumbar enlargement. Although, only for section 7 was this difference significant (P= 0.0079). B. Epifluorescent image showing a cluster of ChAT immunoreactive motoneurones (green) some of which show cleaved caspase-3 (red) immunoreactivity in the ventral horn of a G127X mouse. Scale bar 30μm. C. Epifluorescent images showing immunohistochemical labelling of a spinal section taken with a 5x objective to show the location of the cells in the transverse plane. Highlighted is a ChAT immunoreactive neurone (green) close to the central canal. Inset is the same ChAT labelling with the cleaved caspase 3 labelling taken with a 20x objective in a single focal plane showing immunoreactivity for cleaved caspase-3 in red. Scale bar 20 μm. D. Epifluorescent image showing ChAT immunoreactive neurones (green) in a rostral transverse spinal section. In this, labelling of cells in the intermedio-lateral nucleus and intercalated nucleus can be seen. Inset show the highlighted neurones imaged which a 20 × objective in a single focal plane with one cell showing immunoreactivity for cleaved caspase-3 (arrow, red). Scale bar 50 μm.

## DISCUSSION

AISs of spinal motoneurones in the G127X SOD1 mouse model of ALS increase in length at symptom onset. Assuming that the density of Nav 1.6 stays uniform, this would result in an overall increase in the amount of Na^+^ channels at the AIS. This fits with the results of our electrophysiological experiments showing an increase in the rate of rise of the AIS action potential in spinal motoneurones from G127X mice. A constriction of the AIS width could also condense the Na^+^ channels in this region, reducing the load capacitance and increasing the excitability of the AIS. An increase in Na^+^ channels at the AIS, along with increased activation of I_h_ currents and the earlier onset of the secondary range (indicative of increased persistent L-type Ca^2+^ channels) all point to a sudden increased excitability of spinal motoneurones at symptom onset.

### No reductions in the ability to fire repetitively but evidence for a loss of fast motor units

Here, for the first time we demonstrate *in vivo* intracellular recordings in ALS mice at a clear symptomatic stage. Surprisingly, the motoneurones seemed remarkably healthy showing no deficits in the ability to initiate repetitive firing. Nor were there any significant differences in the input-output gain in the primary range. These data suggest that discrepancies in results from the previous experiments in the G127X mice (Bonnevie et al., 2020) and in the G93A mice (Delestree et al., 2014) with respect to repetitive firing cannot simply be explained by differences in disease stage as the mice in the current experiment were clearly symptomatic, with hind-limb motor deficits and muscle atrophy. One caveat with this conclusion is that the motoneurones in the current experiments were identified by antidromic identification following stimulation of the peripheral axon. We have previously shown a progressive disruption of nodes of Ranvier in the ventral roots in this model, which already becomes evident at presymptomatic time points, suggesting that axon degeneration precedes cell loss in this model, consistent with a ‘dying-back’ theory (Maglemose et al., 2017). It is, therefore, possible that the motoneurones first affected may not be conducting antidromically. If we are to assume that all remaining motoneurones were conducting antidromically then cell loss with a selective survival of cells with longer AISs is unlikely to explain our results as the cell counting revealed surprisingly minimal cell loss in this model.

The ability to repetitively fire and a similar I-f gain within the functional working range for tetanic contractions (Jensen et al., 2018) between the G127X mice and WT suggested that the neurones from which we recorded in the G127X mice were still functional, despite a clear reduction in overall motor function. It is possible that plastic changes at the AIS along with increases in I_h_ currents could represent a mechanism to maintain this function. Nav 1.6 mediates both fast transient and slow persistent inward currents. The latter are important to kick off repetitive firing, especially under conditions of reduced input (Li and Bennett, 2007) and have been indirectly shown to be increased in the G93A mouse (Delestree et al., 2014). A similar trend was observed in the current study in the G127X (although a low ‘n’ in the G127X group most likely affected statistical power). If increases in PICs are an important mechanism to maintain repetitive firing in response to an increase in cell size as have been proposed (Delestree et al., 2014), then motoneurones in these mice would be more sensitive to the use of anaesthesia, in particular the barbiturate anaesthesia used in the G93A experiments, that have been shown to block PICs (Button et al., 2006;Guertin and Hounsgaard, 1999). This could potentially explain the differences in repetitive firing abilities observed between the previous experiments in the G127X and the G93A mice.

The one deficit observed was a reduction in the maximum firing frequencies seen in the G127X mice. From the distribution of the individual data points for maximum firing frequencies in the G127X mice (Fig. 5H), it can be hypothesised that the difference could simply be a loss of the motoneurones with the fastest firing frequencies rather than a loss of ability of individual motoneurones to fire at high frequencies. It has been well documented in other SOD1 models that fast-fatigable motoneurones are the most vulnerable subtype of motoneurone in this disease (Bradley et al., 1983;Dengler et al., 1990;Fischer et al., 2004;Hegedus et al., 2007;Hegedus et al., 2008;Valdez et al., 2012). This reduction in maximum firing frequencies could also, however be affected by many other features (such as increased fatigue or spike frequency adaptation) as the maximum firing frequencies were measured at the peak of current ramps.

Detachment of motoneurones from their muscles fibres is an early change observed in ALS (Moloney et al., 2014) and may also contribute to AIS changes and altered excitability. Detachment of motoneurones from the their targets, either physically through axotomy or functionally using toxins that block neurotransmission has also been shown to alter central motoneurone excitability (Foehring et al., 1986;Kuno and Llinas, 1970;Nakanishi et al., 2005;Pinter and Vanden Noven, 1989;Pinter et al., 1991) thus, it can plausibly contribute to the changes we observed. In particular, blocking of synaptic transmission at the neuromuscular junction using botulinum toxin cause increases in AIS length, albeit of lower magnitude than observed here (Jensen et al., 2020).

### Increases in AIS Na^+^ channels/currents represent a plastic change at symptom onset

Fasciculations, cramps, hyperreflexia and spasticity are symptoms observed in ALS patients and are consistent with an increased excitability of lower motoneurones. Threshold tracking techniques have offered a way to investigate the excitability of the more peripheral motor axons in ALS patients. Such investigations have generally found an imbalance of K^+^ and Na^+^ currents, in particular, increases in persistent Na^+^ currents (Bostock et al., 1995;Cheah et al., 2012;Nakata et al., 2006;Shibuta et al., 2010;Shibuya et al., 2013;Stephanova et al., 2012;Vucic and Kiernan, 2010). This technique has also been applied to the peripheral motor axons of G127X mice, which also showed signs consistent with increased persistent Na^+^ currents (Moldovan et al., 2012). The results of the current study show on the single cell level that the increased Na^+^ currents are not restricted to the peripheral axons but are also a feature of the proximal axon, most importantly, at the site of action potential initiation.

The increased AIS length that we have observed, along with the changes in Na^+^ channel activity at the AIS, would appear to represent plastic changes that occur around symptom onset as they are not seen in presymptomatic mice, approximately 1 month younger (Bonnevie et al., 2020). In fact, in the former study the AISs were actually 6.5% shorter than WT so it can be concluded that the overall increase in length at symptom onset may be even larger, which is in contrast to the tendency towards AIS shortening we have observed on motoneurones in middle age (unpublished observation).

### Does increased excitability contribute to excitotoxic cell death?

The only established treatment for ALS patients that has been documented to prolong survival is Riluzole, which only extends survival by approximately 3 months (Miller et al., 1996;Miller et al., 2007). A new drug (Radicava) has recently received FDA approval aimed at prolonging survival by targeting oxidative stress. The precise mechanisms by which Riluzole prolongs survival are unclear. In addition to interfering with glutamatergic transmission (Azbill et al., 2000;Dunlop et al., 2003;Wang et al., 2004) in animal models, Riluzole has been shown to also block both inactivated Na^+^ channels (Benoit and Escande, 1991) and slow persistent Na^+^ currents (Urbani and Belluzzi, 2000). At clinical dosages in ALS patients, Riluzole has been shown to both normalise the cortical hyperexcitability seen in these patients (by restoring deficits in short interval intracortical inhibition) and, importantly, to reduce peripheral axonal Na^+^ currents of the lower motoneurones, with a stronger effect on inactivated voltage gated Na^+^ channels than persistent Na^+^ conductances (Geevasinga et al., 2016;Vucic et al., 2013). This further suggests that increased Na^+^ currents play an active role in the progression of the disease.

Due to the high levels of Ca^2+^ entering the cell during an action potential, changes at the AIS (being the site of action potential generation) would have potentially large consequences for Ca^2+^ -mediated excitotoxicity. This would especially be a problem when combined with the low Ca^2+^ buffering ability of motoneurones, combined with mitochondrial and astrocytic dysfunction which are also observed in SOD1 mutants (Bowling et al., 1993;Carri et al., 2016;Das and Svendsen, 2015;De Vos et al., 2007;Diaz-Amarilla et al., 2011;Fritz et al., 2013;Haidet-Phillips et al., 2011;Phatnani et al., 2013;Rojas et al., 2014;Shi et al., 2010). However, disruptions of nodal organization consistent with axon degeneration are already occurring in presymptomatic G127X mice, one month younger than in the current study, when the AIS is actually shorter (Bonnevie et al., 2020). It is therefore unlikely that increased AIS Na^+^ currents are initiating the distal degenerative process, although it is still possible that they may contribute to the subsequent cell death. This may explain the limited effects of Riluzole in humans and also why it has been shown in transgenic mouse models of the disease to slow disease progression but not prevent or delay disease onset (Gurney et al., 1996).

Increased I_h_ currents, indicated by increased sag amplitude have also been observed in pyramidal neurons in layer 5b of the cortex of G93A SOD1 mice along with a higher expression of HCN channel genes in the motor cortex compared to controls (Buskila et al., 2019). Interestingly, it was hypothesized that I_h_ could also be enhanced by a reduced activity of Kv7 channels which would fit well with observation from induced pluripotent stem cells derived from ALS patients showing a reduction in delayed-rectifier K^+^ currents. In those experiments a specific activator of subthreshold Kv7 currents corrected a hyper-excitable phenotype (Wainger et al., 2014). Our results suggest that this alteration in I_h_ currents is not restricted to the motor cortex but also seen in the lower spinal motoneurone in the spinal cord of ALS mice *in vivo*.

A potential excitotoxic role for the increased excitability at this stage would be consistent with a lack of obvious cell death immediately prior to symptom onset in this model (Bonnevie et al., 2020) and the large amount of motoneurones now expressing cleaved caspase-3 in the symptomatic mice. The cleaved caspase-3 labelling also confirms that cell death in this model is via apoptotic pathways as has been shown in other SOD1 models (Pasinelli and Brown, 2006). Our observation of cleaved caspase-3 activity in ChAT immunoreactive neurones other than motoneurones is also consistent with growing evidence that, although motoneurones are particular vulnerable in this disorder, they are not exclusively affected. The observed cleaved-caspase-3 activity in neurones in the IML/intercalated nucleus is consistent with similar observations in the G93A mice with a loss of IML neurones (Kandinov et al., 2013) which is also observed post-mortem in human ALS patients (Hayashi et al., 2016;Shimizu et al., 2000;Takahashi et al., 1993). Consistent with this, there is now considerable evidence for autonomic abnormalities in both SOD1 mouse models (Kandinov et al., 2011) and ALS patients (reviewed by (Fang et al., 2017)).

Cleaved caspase-3 activity in ChAT-immunoreactive neurones around the central canal (lamina X) is of particular interest, given that this population of cells includes those giving rise to the C-boutons observed on alpha motoneurones (Zagoraiou et al., 2009). A loss of these cells has been shown in G93A mice (Milan et al., 2015), and changes in C-boutons frequency or size have also been observed (Herron and Miles, 2012;Milan et al., 2015;Pullen and Athanasiou, 2009;Saxena et al., 2013) although the results have been inconsistent, which may be due to experimental design (Dukkipati et al., 2017).

The hypothesis of an increased motoneurone excitability contributing to excitotoxic cell death in this disease has been questioned by a study that attempted to enhance motoneurone excitability. This intervention actually promoted neuroprotection of motoneurones (Saxena et al., 2013). The important caveat with those experiments, however, is that the final intrinsic excitability of the motoneurone was never tested. This would have been crucial as it is entirely plausible that the methods used to increase excitability would have been just as likely to drive homeostatic mechanisms, potentially reversing the AIS plasticity that we have observed here.

### Increases in AIS sodium channels/currents may contribute to spasticity

The appearance of an increased excitability around symptom onset suggests that such changes could help to explain some of the gains of function observed in the disease, in particular fasiculations, hyperreflexia and spasticity. In models of spinal cord injury it is now well documented that hyperreflexia and spasticity appear as a consequence of increased motoneurone excitability in addition to local changes in spinal cord circuitry. This includes a reduction of inhibition (Norton et al., 2008), reductions in post-activation depression of excitatory Ia afferent synapses on motoneurones (Schindler-Ivens and Shields, 2000), increases in Na^+^ and Ca^2+^ PICs (Bellardita et al., 2017;Eken et al., 1989;Murray et al., 2010), and an upregulation or constitutive activity of serotonergic and noradrenergic receptors (Rank et al., 2011;Murray et al., 2010).

Spasticity is also observed in SOD1 models of ALS (Dentel et al., 2013;Modol et al., 2014), and many of the changes observed following spinal cord injury have also been observed in SOD1 models, including the G127X. Reductions in post-activation depression have been observed in the G127X mouse in both presymptomatic and symptomatic mice (Hedegaard et al., 2015). Na^+^ and Ca^2+^ PICs are increased presymptomatically in the G93A (Delestree et al., 2014) and G127X models (Bonnevie et al., 2020;Meehan et al., 2010) respectively and in symptomatic G127X mice (reported in the current experiments). Given that these changes are seen in presymptomatic mice, it would appear that they alone cannot account for the later symptoms such as spasticity. In the symptomatic period, however, these changes would be exacerbated by an upregulation of 5HT2C receptors, which also show constitutive activity (Dentel et al., 2013). In addition would be reductions in the recurrent inhibition of spinal motoneurones due to a loss of Renshaw cells (Chang and Martin, 2009;Chang and Martin, 2011b;Chang and Martin, 2011a). Finally, there is the contribution from the plastic changes in the motoneurone AISs that we have shown here for the alpha motoneurones and potentially also occurring in gamma motoneurones (that control the sensitivity of muscle spindles). We propose that it is a combination of all these factors that interact to produce the hyperreflexia and spasticity in ALS.

### Implications of our results for potential new treatments

Our results provide the first definitive *in vivo* evidence of how central lower motoneurone excitability increases as the disease progresses in this mouse model of ALS. The results of our work suggest that targeting AIS plasticity may provide a therapeutic strategy to not only target disease progression, but also as a symptomatic treatment for spasticity and hyperreflexias in ALS. Blocking Na^+^ channels per se risks reducing the residual function of already weak muscles. Targeting the mechanisms underlying the AIS plasticity may, however, provide a more optimal strategy. Here, our observations of an earlier activation for the secondary range of firing is important as the secondary range is believed to be due to the delayed activation of dendritic L-type Ca^2+^ channels, when direct current injection to the soma is used to mimic synaptic input. This is relevant as activation of L-type Ca^2+^ channels has been shown in other neurones to mediate activity dependent plasticity of AISs (Evans et al., 2013;Grubb and Burrone, 2010). It is therefore possible that the changes in AISs that we have observed are driven by altered L-type Ca^2+^ channel activation. Targeting L-type Ca^2+^ channels or more downstream signalling cascades in spinal motoneurones could potentially therefore provide an optimal therapeutic target for spasticity by blocking AIS plasticity.

## Acknowledgments

We thank Karin Graffmo & Stefan Marklund, Umea University, for originally providing the G127X SOD1 mouse line. We thank Lasse Luab and Niels Esben for ChAT labelling of one of the symptomatic mice used for cell counting. All confocal analysis was performed at Core Facility for Integrated Microscopy (CFIM) at the University of Copenhagen, Faculty of Health and we thank their staff for their unfailing help and advice.

---

**Supplementary Table 1.**
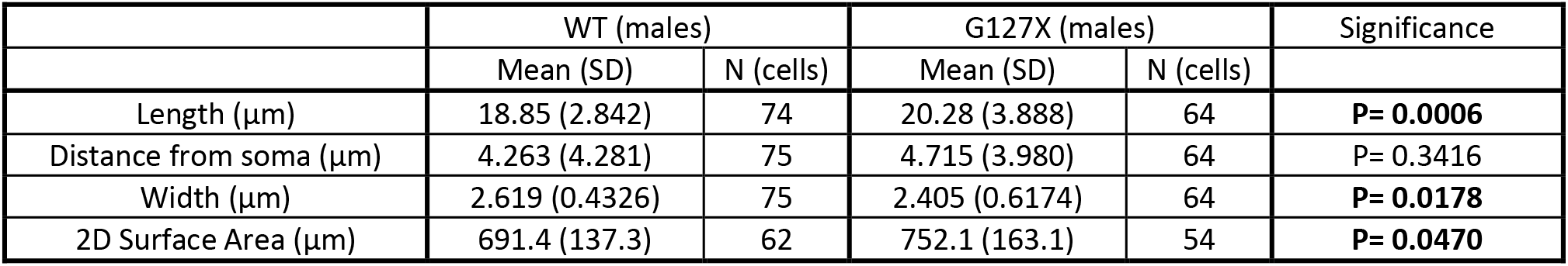
AIS parameters with only males in both groups.

**Supplemental Table 2.** AIS parameters of gamma motoneurones from the mixed sex groups.

|  | WT |  | G127X |  | Significance |
| --- | --- | --- | --- | --- | --- |
|  | Mean (SD) | N (cells) | Mean (SD) | N (cells) |  |
| Length ($\mu\text{m}$) | 17.57 (4.068) | 34 | 19.57 (3.525) | 21 | P = 0.0684 |
| Distance from soma ($\mu\text{m}$) | 5.194 (3.681) | 34 | 5.788 (4.944) | 21 | P = 0.9350 |
| Width ($\mu\text{m}$) | 1.950 (0.7231) | 34 | 1.979 (0.5645) | 21 | P = 0.8773 |
| 2D Surface Area ($\mu\text{m}$) | 344.4 (97.17) | 34 | 380.7 (91.21) | 21 | P = 0.1749 |

